# Perturbation of epithelial and limbal stem cell identity in a mouse model of pathologic corneal neovascularization

**DOI:** 10.1101/2025.05.22.655502

**Authors:** Mojdeh Abbasi, Julian A. Arts, Dina Javidjam, Petros Moustardas, Ava Dashti, Daniel Aberdam, Huiqing Zhou, Neil Lagali

**Author notes:** equal contributions (Mojdeh Abbasi, Julian A. Arts). senior authors (Huiqing Zhou, Neil Lagali). This work was performed at Linköping University, Linköping, Sweden and at Radboud University, Nijmegen, The Netherlands. **Correspondence should be addressed to:** Neil Lagali, Department of Biomedical and Clinical Sciences, Linköping University, SE-581 83 Linköping, Sweden; Jo Huiqing Zhou, Department of Molecular Developmental Biology, Radboud University, 6525 GA Nijmegen, The Netherlands.

## Abstract

The epithelial layer of the cornea is a critical physical and ocular immune barrier for maintaining tissue integrity, homeostasis, and transparency for proper vision. Corneal injury can trigger inflammation, impair wound healing and compromise immune privilege and avascularity, leading to vision loss. Moreover, injury to the cornea can disrupt the engine of epithelial repair and restoration, the limbal stem cell (LSC) niche. Here we used a corneal suture model to induce epithelial damage, sustained inflammation and neovascularization, to examine the impact on LSCs. Using single-cell transcriptomics, we analyzed corneal cell state changes and additionally evaluated the potential of duloxetine, an FDA-approved medicine, to promote wound healing and corneal homeostasis. Single-cell RNA-seq analysis revealed loss of homeostatic limbal stem cells, basal and differentiated epithelial cells and an increase in distinct limbal-like, conjunctival, inflammatory, and vascular cell states, suggesting a coordinated wound healing response in different tissue layers. Importantly, duloxetine treatment promoted epithelial homeostasis, enhanced stem cell-like and stromal repair processes, and suppressed immune and vascular responses. Examination of corneal cell perturbation and transformations at the single-cell level thorough marker profile annotations can improve the understanding of LSC plasticity and function while yielding potential biomarkers of corneal repair processes.

## Introduction

The cornea is the eye’s transparent outermost tissue and its first-line mechanical and immune barrier. The corneal epithelium in particular serves a vital role for maintaining corneal integrity and homeostasis, which is essential for maintaining the cornea’s transparency. Injury or diseases of the cornea can lead to inflammation and loss of both corneal immune privilege and avascularity^1^. This is a consequence of disturbing a complex balance of epithelial and stromal cytokines and pro- and anti-angiogenic factors^1–3^. In particular, a functioning limbal stem cell (LSC) niche that provides continuous renewal of a normal corneal epithelium is pivotal for providing this fine balance of factors that prevents the non-transparent, vascularized conjunctival epithelium from migrating into the cornea ^4^.

In conditions characterized by chronic or impaired wound healing, such as in traumatic or congenital limbal stem cell deficiency (LSCD), the cornea loses its immune and angiogenic privilege ^5,6^ combined with the breakdown of the limbal barrier ^4^. The result is inflammation, corneal epithelial transformation and neovascularization, with neovascularization characterized by the aberrant invasion of blood (hemangiogenesis) and lymphatic (lymphangiogenesis) vessels originating from the limbal vascular arcade and conjunctiva into the normally avascular cornea ^7–11^. Consequently, the cornea loses its capacity for proper and efficient wound healing, exacerbating the cycle of damage and repair failure ^12^. What is less appreciated, however, is that chronic inflammation can simultaneously impact the limbal stem cell environment, potentially leading to loss of LSC function that may be temporary or prolonged.

Experimental models for corneal neovascularization, such as alkali burn, mouse injury or transplantation models ^10,13–16^ and corneal suturing ^7,8,10^ are common. Among these, the corneal suture model has gained prominence for its ability to closely mimic the wound conditions and sustained inflammation that drive corneal neovascularization, and for its excellent reproducibility. This model involves placing surgical sutures in the corneal stroma, leading to disruption of the corneal epithelial barrier, introduction of a foreign body (suture material) into the stroma and persistent mechanical irritation by the knotted suture end ^17^. These events collectively initiate an inflammatory wound-healing cascade in the cornea, with subsequent blood vessel invasion, epithelial transformation and loss of corneal transparency ^18^. This sequence of events is observed in many clinical situations, for example in the context of corneal transplant rejection ^19^, alkali burns ^20^, and aniridia-associated keratopathy ^21^.

Although angiogenic processes involved in corneal neovascularization have been studied in the suture model ^7–10^, the contribution of various cell types and structures in the cornea – especially the epithelial barrier, including the limbal stem cells and their niche and their potential disruption upon suturing – is not well studied. In addition, neovascularization is an important component of LSCD in traumatic (chemical burns) or congenital cases like aniridia ^22^, a disease of *PAX6* haploinsufficiency caused by genetic mutations affecting this key transcription factor. *PAX6* controls corneal epithelial identity and maintenance during development and in the postnatal period. Inflammation and neovascularization, common features of LSCD, result in the downregulation or loss of PAX6 protein ^23^ while congenital aniridia, where a reduced PAX6 protein level has been reported, is characterized by progressive LSCD and corneal neovascularization ^24,25^. Enhancing PAX6, therefore, could possibly preserve epithelial identity to attenuate corneal neovascularization and LSCD ^26–28^.

Duloxetine, an FDA-approved medicine for treating severe depression, is a serotonin– norepinephrine reuptake inhibitor known to inhibit phosphorylation of the p38 MAPK (MEK/ERK) pathway thus inhibiting NF-kB nuclear translocation and upregulating PAX6 ^29,30^. By MEK/ERK pathway inhibition, duloxetine was shown to enhance PAX6 protein levels *in vitro* in human telomerase-immortalized CRISPR-Cas9 mutated *PAX6^+/-^* limbal stem cells (mut-LSCs) and thereby promote epithelial wound healing *in vitro* ^29^.

The aim of this study is therefore two-fold - first, to examine the process of corneal neovascularization in the mouse at the single-cell transcriptomic level with particular attention to effects on the limbal stem cell niche; and secondly, to explore whether duloxetine could potentially rescue aberrant wound healing and neovascularization in this context.

## Results

### Corneal sutures induce a sustained epithelial wound triggering inflammation, corneal nerve degeneration and neovascularization

Ten days following intrastromal suture placement in the mouse cornea, slit lamp analysis revealed extensive corneal neovascularization from the limbus towards the central cornea. The vascular response did not differ in clinical appearance in eyes treated for 10 days with duloxetine (treatment group), relative to treatment for 10 days with the PBS vehicle only (suture group, Figure 1A). Laser-scanning *in vivo* confocal microscopy revealed significant neovascular sprouting and a marked increase in inflammatory cell infiltration within the epithelium and corneal anterior-deep stroma relative to the no suture group (p<0.0001), however, no significant difference in inflammatory cell count was noted between treatment and suture groups (Figure 1A, B). Immunofluorescence analysis of corneal flat mounts stained with the neuronal marker β III tubulin revealed pronounced changes in the corneal nerve architecture in suture and treatment groups (Figure. 1C, E). Suturing disrupted the distinct whorl pattern that is characteristic of corneal subbasal nerves and significantly reduced the subbasal nerve density relative to the no suture group (p<0.0001) with the central corneal region most affected (Figure. 1C, D). Similarly, stromal nerve density was significantly reduced following suturing (p<0.0001, Figure. 1E, F). For both subbasal and stromal nerves, however, no differences were noted between suture and treatment groups. The neovascular response was quantified by CD-31 immunostaining, with a significant increase in vascular density noted in the suture group relative to the no suture group (p<0.0001), which was not reduced by treatment (Figure. 1G, H). In summary, this acute and aggressive suture model induced a sustained epithelial wound leading to inflammatory, neurodegenerative and angiogenic responses at the tissue level that were not ameliorated with treatment.

**Figure 1.**
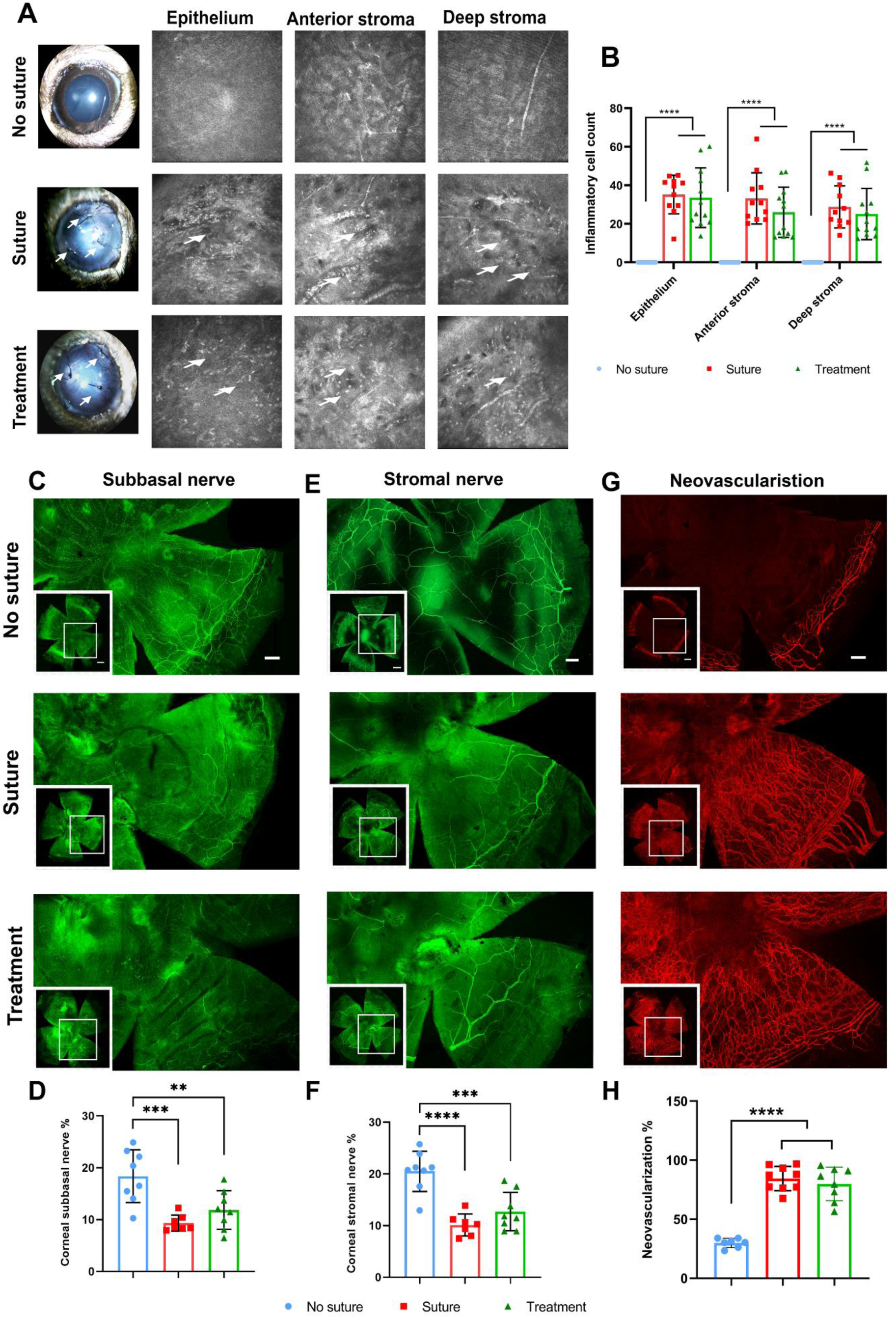
Cellular and phenotypic aspects of corneal neovascularization. (A) Slit lamp photographs depicting loss of corneal transparency and vessel ingrowth from the limbus extending into the cornea following placement of three sutures. (B) *In vivo* confocal microscopic imaging revealed inflammatory cell infiltration (arrows) in different layers of the cornea and quantitative analysis revealed significant increase in inflammatory cells in all layers of sutured corneas relative to non-sutured corneas. (C, D) Representative images of whole-mount βIII tubulin immunostaining of the corneal subbasal and (E, F) stromal nerves and (G, H) Representative corneal wholemounts indicating hemangiogenesis by CD31 staining. Images are magnified views of the region indicated in the inset on the lower left corner. Scale bars denote 500 µm in low magnification images (inserts) and 200 µm in the higher magnification main images, and apply equally to all three vertically aligned images. The data in (B), (D), (F), and (H) are presented as mean ± SD; One-Way ANOVA

### Single-cell RNA-seq analysis identifies profound cornea cell type transformations after suture injury

Single-cell RNA sequencing (scRNA-seq) was performed on corneas retrieved from all groups (Figure 2A). In total 4114 cells for no suture, 3813 cells for suture and 5277 cells for treatment groups were identified within QC thresholds and further analyzed. To identify cell states in each experimental condition, we used Leiden clustering ^31^ on data from each experimental group to unbiasedly classify cells based on their expression profiles (Figures 2B-D). Subsequently, clusters with similar expression patterns of selected marker genes from each experimental group were joined into a ‘cell state’ (Figure S1). The number of distinct cell states increased in the suture and treatment groups, with patterns of marker expression distinct from the no suture group.

**Figure 2:**
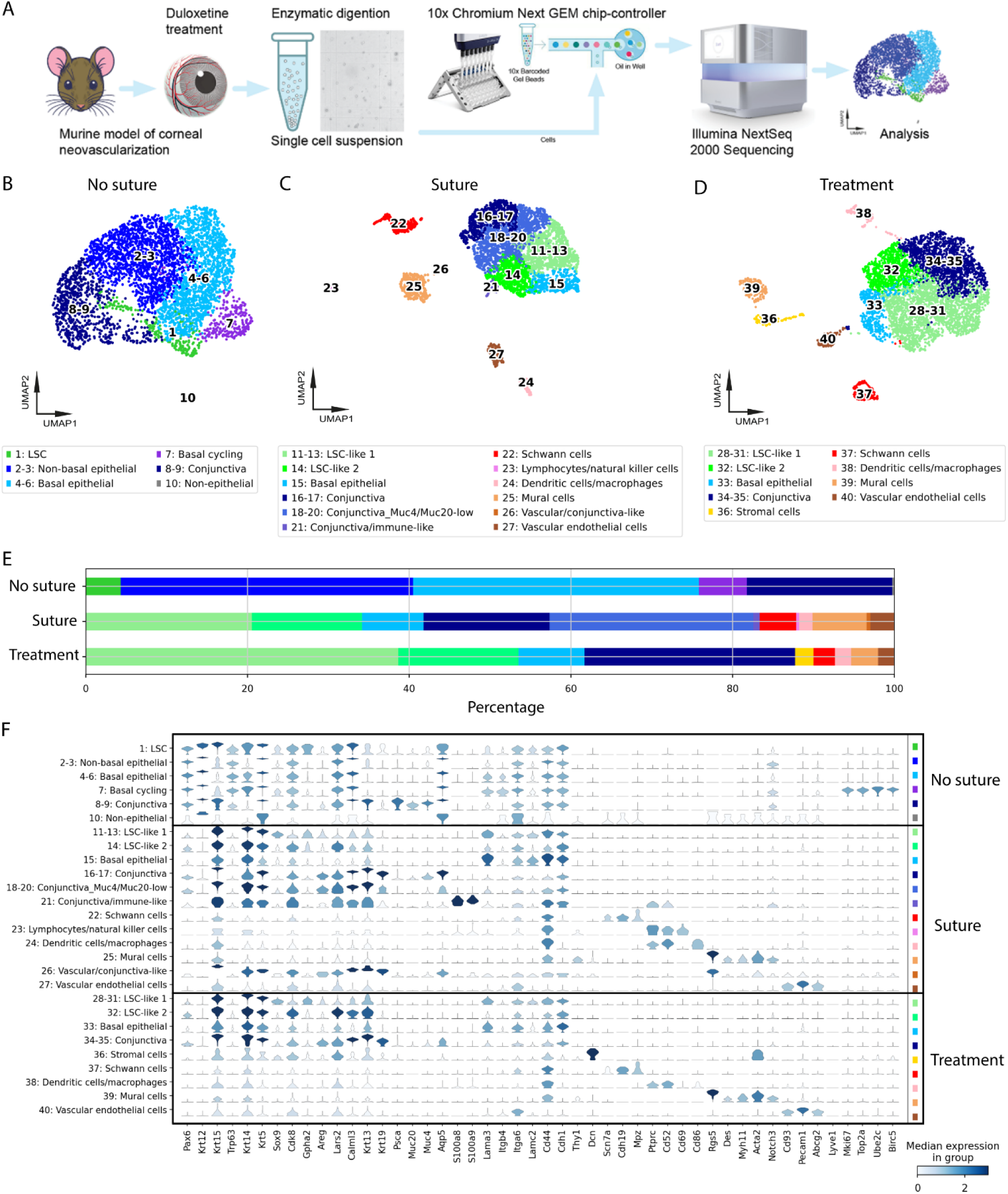
Identification and characterization of cell states in uninjured, sutured and Duloxetine treated corneas. (A) Schematic representation of the experimental strategy and workflow overview in this study. The diagram illustrates the use of a suture model in mice corneas, where treated with duloxetine/ vehicle groups were subjected to downstream single-cell analysis. (B-D) UMAPs of annotated cell states across conditions: no suture (B), suture (C) and treatment (D). (E) Bar plots of relative cell state contributions across conditions. Exact percentages are given in Table S1. (F) Violin plot of marker gene expression across conditions. Similar colors define similar cell states across conditions.

Additionally, we performed integrated analysis with scVI ^32^ using pooled data from the three experimental groups (Figure S2), confirming all cell states identified by the clustering analysis of each individual group. Furthermore, this analysis showed that cells from the suture and treatment groups had gene expression patterns that were similar, while both were distinct from the no suture group.

### Cell states in the non-sutured cornea are primarily epithelial

In the no suture group, 6 distinct cell states (10 clusters) were identified. The overwhelming majority of cells (>99%) were characterized as epithelial in nature, classified into 5 cell states (9 clusters). Only very few non-epithelial cells were identified, and these were classified as a single ‘non-epithelial’ cell state (1 cluster).

The cell state LSC, comprised of cells in cluster 1 (5% of the cells), was recognized by expression of limbal markers such as Pax6, Trp63, and high expression of Krt15 and the LSC progenitor marker Gpha2 (Figure 2F) ^33^. The corneal (non-basal) epithelial cell state (clusters 2-3; 30% of the cells) was annotated by high expression of Pax6, a moderate level of Trp63, and a low level of Krt14 (Figures 2E and 2F) ^34^, whereas a basal epithelial cell state (clusters 4-6; 32% of the cells) was characterized by high expression of Trp63 and Krt14 as well as basal markers Lama3 and Itgb4 (Figure 2F) ^35^. A basal cycling cell state (cluster 7; 7% of the cells) showed presence of cell cycle markers such as Mki67, Top2a, Ube2c, and Birc5 ^36^, while still expressing basal epithelial markers comparable to the basal epithelial cell state (Figure 2F). A conjunctival cell state (clusters 8-9), which accounted for around 20% of the total cells, was identified based on expression of conjunctival markers including Krt13, Psca, Muc20, Muc4, and Aqp5 (Figure 2F)^37^. Finally, the smallest cell state (cluster 10; <1% of the cells) clustered away from all other cell states, and had low, but unique expression of non-epithelial markers such as Scn7a, Cdh19, Mpz, Cd93, and Pecam1, and Abcg2 (Figure 2F). Integrated analysis showed that this cluster consisted of an extremely low number of vascular endothelial cells, mural cells and Schwann cells (Figure S2B).

### Suturing dramatically alters homeostatic corneal epithelial cell states

Globally, cell states were significantly altered in the suture group, as compared to the no suture group (Figure 2C & 2E). In total, 12 cell states, composed of 17 different clusters, were identified. Among them, 3 limbal and corneal epithelial related cell states, 3 conjunctival cell states, and 6 non-epithelial cell states, which represented a 17% increase in the number of non-epithelial cell states.

In the suture group, the typical LSC cell state with high Pax6, Trp63 and Gpha2 expression, making up 5% of the cells in the non-suture group, was not detected. Instead, cell states with high expression of the limbal marker Gpha2 but reduced expression of Pax6, Trp63, Krt5, Krt19, and Aqp5, as compared to LSCs in the no suture group (Figure 2C and Figure 2F), were detected (clusters 11-14). These cell states accounted for more than 34% of the cells in the suture group, a significant increase of Gpha2 positive cells, and were thus annotated as ‘LSC-like’. The reduced Pax6 and increased Gpha2 expression at the limbal region was validated at the protein level (Figure 3A and 3B). The LSC-like cells were further divided into two sub-cell states, LSC-like 1 (cluster 11-13) because of high expression of Sox9 and low expression of Cdk8, as well as relatively high expression of Areg and low expression of Lars2, and into LSC-like 2 (cluster 14) because of the inverse expression pattern (Sox9 (low) and Cdk8 (high), Areg (low) and Lars2 (high)) of these genes (Figure S1).

**Figure 3:**
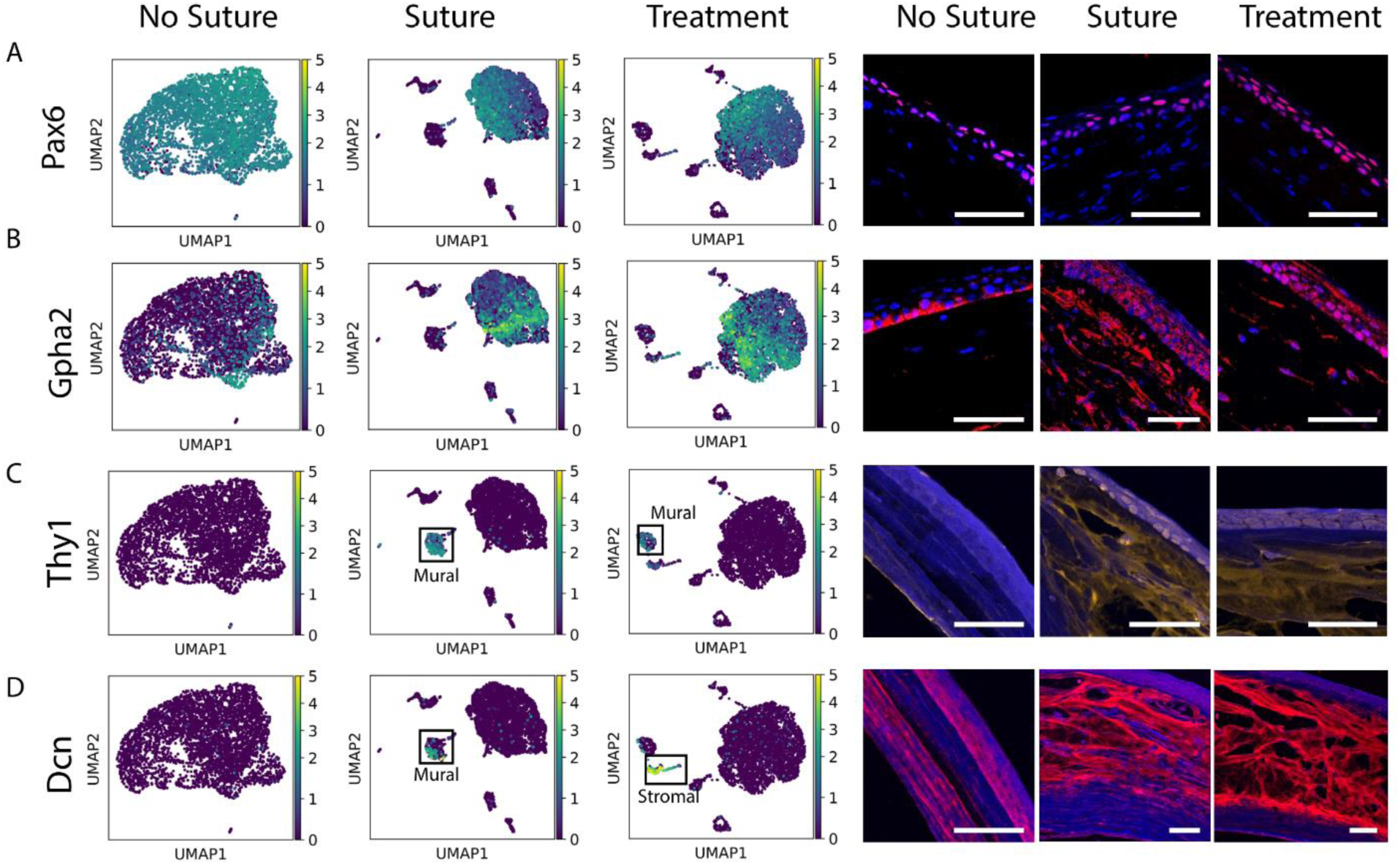
Gene expression and protein staining of affected marker genes in sutured corneas. Expression and staining respectively of gene-protein targets detected in single-cell RNA-seq data shown by UMAPs and at the protein levels shown by immunohistochemistry staining for Pax6 (A), Gpha2 (B), Thy1 (C) and Dcn (D). Nuclei are stained with DAPI. Scale bars represent 50 µm in all figures.

Suturing also drastically modified other corneal epithelial cells. Together, the corneal epithelial cell states (clusters 2-7) declined by 69% in the suture group relative to the no suture group (Figure 2E, Table S1). Strikingly, basal cycling cells were completely absent in the suture group, as none of the four cycling markers identified in the no suture group were present in the suture group (Figure 2F). Additionally, non-basal epithelial cells with high Pax6 but with low Krt14 expression were completely absent in the suture group. Only one corneal epithelial cell state expressing the basal epithelial markers Krt15, Krt14, Krt5, Lama3 and Itga6 (cluster 15) remained in the suture group, and was therefore annotated as basal epithelial. But this epithelial cell state had relatively low expression of Itgb4, Pax6 and Trp63, as well as an increase in Krt16 and Krt13, relative to basal epithelial cells in the no suture group, highlighting the altered cell expression pattern of basal epithelial cells compared to their healthy counterparts (Figure 2F, and 4B).

### Suturing promotes the emergence of conjunctival, inflammatory and vascular cell states

In contrast to the single distinct conjunctival cell state in the non-suture group, 3 conjunctival related cell states were detected in the suture group. Two relatively normal conjunctival cell states (clusters 16-20) expressing Krt13, Krt19, Muc4, and Aqp5 were annotated, showing a 24% increase in total cell population in the suture group, relative to the no suture group (Figure 3D). These cell states were separated into 2 sub-cell states, as one cell state (clusters 18-20) had significantly lower expression levels of well-known conjunctival markers Muc4 and Muc20 than the other cell state (clusters 16 and 17). An additional conjunctival related cell state (cluster 21; <1% of the cells) was identified specific to the suture group, expressing both conjunctiva markers and inflammatory mediator genes S100a8 and S100a9 (Figure 2F) ^38,39^. This cell state was therefore annotated as conjunctiva/immune-like. The prevalence of conjunctival cell markers in the corneal tissue upon suturing was confirmed at the protein level by immunostaining for Muc5ac, a canonical conjunctival marker widely used as a conjunctival/goblet cell marker in corneal pathology and LSCD ^40,41^. Muc5ac protein was not present in the no suture group, but was clearly expressed in the epithelium of the sutured groups (Figure S3), indicating conjunctivalization of the superficial epithelium in the sutured groups.

Notably, several non-epithelial cell states emerged in the suture group, which were absent in the no suture group. A non-epithelial cell state (cluster 22; 5% of cells) expressed the markers Scn7a, Cdh19 and Mpz (Figure 3E) and was annotated as Schwann cells ^42^. Two cell states (clusters 23 and 24; 2% of the cells) expressed specific immune cell markers for lymphocytes/natural killer (NK) cells and dendritic cells/macrophages, respectively. Whereas both of these immune cell states expressed Ptprc and Cd52, the lymphocytes/natural killer (NK) cell state uniquely expressed the lymphocyte marker Cd69 ^43^, while the dendritic cells/macrophage cell state expressed the dendritic cell marker Cd86 ^44^ (Figure 3E). The largest non-epithelial cell state (cluster 25; 7% of cells) was annotated as the mural cell state, expressing mural markers Rgs5, Des, and Myh11 ^45^ and Thy1 (Figure 3E). Thy1, a typical mural cell marker ^45^ that was absent in the no suture group and upregulated in the suture group (cluster 25) was in the tissue similarly found to be expressed in a small population of cells in the stroma in the suture group, consistent with neovascularization ^46^. The cell state annotated as ‘mural cells’, however, expressed the markers Myh11, Acta2, Thy1 and Notch3, which are also known fibroblast/myofibroblast markers ^45,47,48^. An additional non-epithelial cell state (cluster 26; <1% of cells) specific to the suture group was annotated as vascular/conjunctiva-like, as cells simultaneously expressed conjunctival markers (Krt13, Krt19, Aqp5) and low expression of Cd93 and Pecam1 (Figure 3E). A vascular endothelial cell state (cluster 27; 3% of cells) was also identified by high expression of vessel markers (Cd93, Pecam1) and expression of Abcg2 ^49^ (Figure 3E).

### Duloxetine treatment suppresses immune and vascular responses while promoting stromal healing and basal epithelial normalization

In the treatment group where Duloxetine was applied, 9 cell states were identified, composed of 13 distinct cell clusters. Three limbal and corneal epithelial cell states, 1 conjunctival state and 5 non-epithelial states were identified.

LSC-like 1 (clusters 28-31; 39% of cells) and LSC-like 2 (cluster 32; 14% of cells), the 2 cell states similar to those detected in the suture group, were present in the treatment group. Nevertheless, in the treatment group, the population of LSC-like 1 cells almost doubled in proportion, from 21% in the suture group to 39% in the treatment group (Figure 2E). Importantly, marker genes for the limbal region *Pax6* and *Ghpa2* exhibited partial rescue of altered expression with treatment at the protein level, with recovery of broader epithelial expression of PAX6 and suppression of stromal expression of GPHA2 (Figure 3A, 3B). While the proportion of basal epithelial cells (cluster 33; 8% of cells) did not change with treatment, these cells nonetheless exhibited similar expression levels of Lama3, Lars2, Lamc2 and Cd44 with treatment, as compared to those observed in the no suture group. Interestingly, a new ‘stromal’ cell state emerged uniquely in the treatment group (cluster 36; 2% of cells), with high expression of Decorin (Dcn). In scRNA-seq data, while Dcn expression was almost completely absent in no suture mice and low in the suture group (clusters 25-26), duloxetine treatment resulted in a significant enhancement in cells annotated as stromal cells (cluster 36). At the protein level, Dcn was expressed at a low level in the basal epithelium and posterior third of the stroma in the no suture group, at a higher level in the anterior two-thirds of the stroma in the suture group, and at the highest levels in the treatment group in all stromal layers (Figure 3D). Furthermore, lymphocytes/natural killer cells (cluster 23) and vascular/conjunctiva-like (cluster 26) cell states which were induced in the suture group were completely absent in the treatment group (Figure 2D). Treatment also suppressed the conjunctiva/immune-like cell state (cluster 21), with a dramatic reduction in S100a8/a9 to the level of the no suture group (Figure 2F). Reduction in the conjunctival cell states with treatment was confirmed by immunostaining of the corneal tissue, where treatment suppressed the expression of the Muc5ac marker (Figure S3). Dendritic cells/macrophages (cluster 38; 2% of cells) remained in the treatment group.

Several non-epithelial states were similarly identified in the treatment group and in the suture group, such as Schwann cells (cluster 37), mural cells (cluster 39) and vascular endothelial (cluster 40); however, the proportion of these cells was reduced by 2%, 4% and 1% respectively, in the treatment group relative to the suture group. In particular, both vascular endothelial cells (from 3% to 2%) and mural cells/fibroblasts (from 7% to 3%) were reduced with treatment. Correspondingly, Thy-1 expression was suppressed in the treatment group detected in scRNA-seq data and confirmed at the protein level (Figure 3C).

### Duloxetine treatment partially suppressed abnormal immune responses and non-epithelial cell states

To further investigate how cell states are affected by suture-induced processes and the potential modulatory effects of duloxetine, we specifically assessed several known pathways that were relevant for the suture conditions based on previous functional characterization and microarray analysis ^10,50,51^ (Figure S4). To do this, we used integrated data of no suture, suture and treatment groups (Figure 2SE) and performed highly variable gene analysis on detected cell states (Table S2). Next, we conducted gene set enrichment analysis (GSEA) on the identified highly variable genes utilizing gene sets in KEGG pathway ^52^ and Gene Ontology Gene ontology enrichment (Ashburner et al., 2000) databases (Table S2). We included the analyzed pathways and biological processes TNF signaling, positive regulation of inflammation, positive regulation of angiogenesis, MAPK signaling given its relevance for duloxetine treatment ^23,29^, and axon guidance given the subbasal nerve degeneration observed in sutured corneas (Figure 1C). Pathways were considered as ‘rescued’ if significance levels decreased after duloxetine treatment relative to the suture group, or if significance levels were restored to levels comparable to the no suture group following duloxetine treatment.

In LSC-like 1 cells (cluster 11-13), highly variable genes showed enrichment for TNF signaling after suturing, but this was suppressed by duloxetine treatment (Figure 4A). This observation was supported by examining the expression of *Tnfrsf1b* (a receptor for TNFα) at the single-cell RNA-seq level (Figure 4B). Staining of the TNFR2 protein, encoded by *Tnfrsf1b*, in the limbus (Figure 4C) showed that the impact of Duloxetine treatment on this protein was substantial, as basal cells of the limbus initially expressed this gene in the no suture group, yet this expression was completely suppressed following duloxetine treatment.

**Figure 4:**
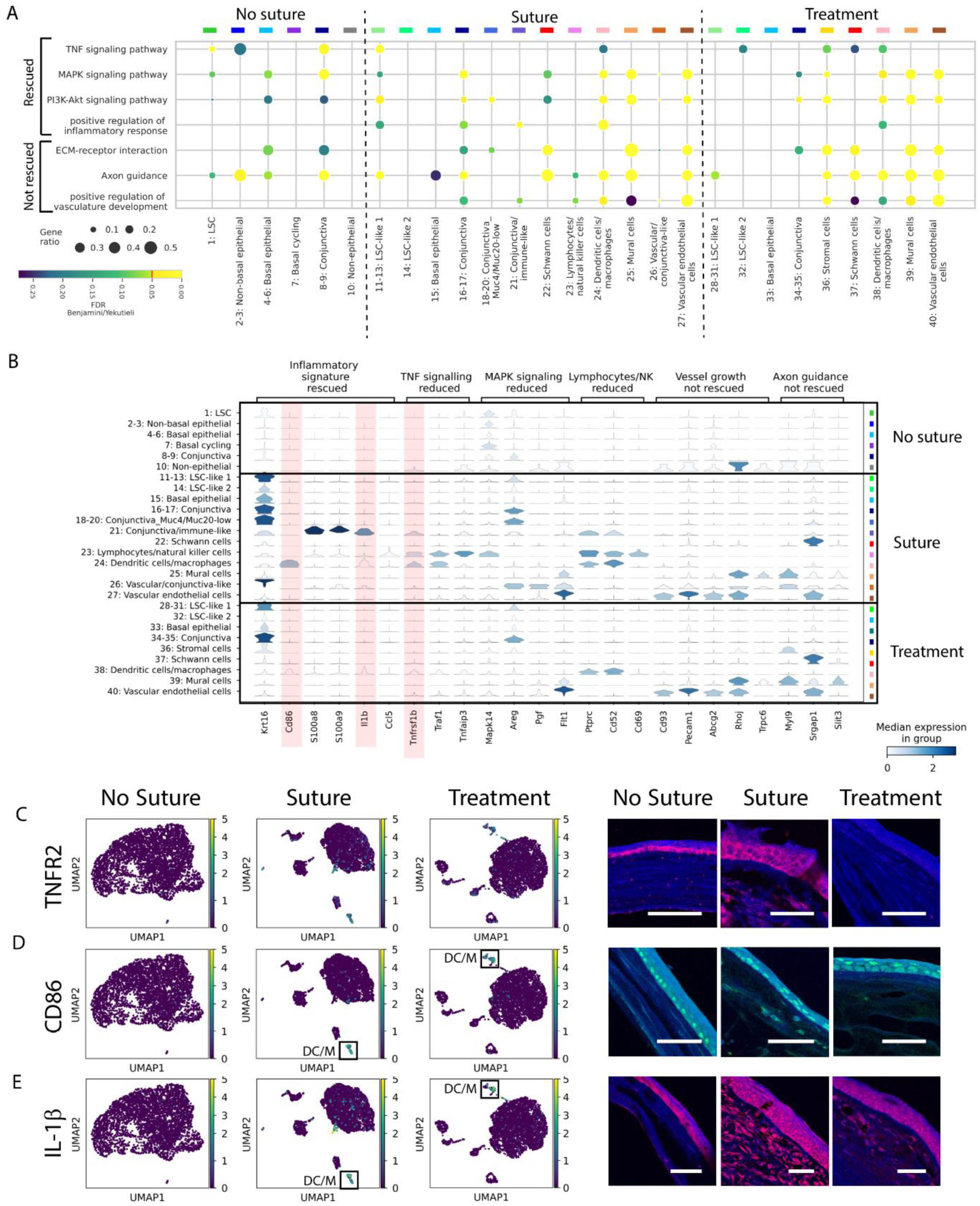
Characterization of disease processes on the single-cell landscape of sutured corneas and effects by duloxetine. (A) Preselected gene sets associated with suture-induced disease processes underwent GO-Term and KEGG pathway enrichment analysis (selected processes from Figure S4) across all cell states within the three experimental groups (no suture, suture and treatment). (B) Violin plots of specific gene expression for suture-induced disease processes. Highlighted genes are stained in lower panels. (C-E) UMAPs and staining for genes in the limbus: TNFR2 (C) and of inflammatory genes in the central cornea: Cd86 (D) and IL-1β (E). The data in (A) are presented as the p-value adjusted using the Benjamini–Yekutieli method. Scale bars represent 50 µm in all figures.

In the suture group, highly variable genes in conjunctival cells were strongly enriched for the MAPK pathway. In addition, genes in several other cell states such as mural cells (cluster 25), vascular/conjunctiva-like cells (cluster 26) and vascular endothelial cells (cluster 27) were also enriched for this pathway. In the treatment group, many of these cell states were lost, and in conjunctiva cells (clusters 34 and 35), enrichment for this pathway was also reduced (Figure 4A). Similar to the MAPK signaling pathway, PI3K-Akt signaling had a comparable enrichment pattern in many cell states, being enhanced in the suture group and suppressed with treatment. The cell states originally showing enrichment for the PI3K-Akt pathway, such as Conjunctiva-Muc4/Muc20-low (cluster 18-20) and Vascular/conjunctiva-like cells (cluster 26) were completely lost with treatment. In addition, LSC-like 1 cells (cluster 11-13) fully lost their enrichment for this pathway after treatment (Figure 4A).

In the suture group, highly variable genes in many cell states, such as LSC-like 1 (cluster 11-13), conjunctiva (cluster 16 and 17), conjunctiva/immune-like (cluster 21) and immune cells (cluster 24), were enriched for inflammation in Gene Ontology analysis. This enrichment was diminished upon duloxetine treatment (Figure 4A). These changes were consistent with the expression of the interleukin-1 beta (IL-1β) and the dendritic and macrophage marker Cd86 ^53^ in scRNA-seq data (Figure 4B) and further confirmed at the protein level in the corneal tissue (Figure 4D). The IL-1β protein level was significantly increased in the sutured corneas and decreased in both limbal and central epithelial cells with Duloxetine treatment (Figure 4D). Cd86 protein was absent in the stroma in the no suture group, expressed by a small population of cells in the stroma in the suture group, and was suppressed by duloxetine treatment (Figure 4E).

Processes such as extracellular matrix (ECM)-receptor interaction, axon guidance and positive regulation of vasculature development remained unaffected following Duloxetine treatment (Figure 4A, Figure S4). Of importance, axon guidance was enriched in highly variable genes of non-epithelial cells (Figure 4A). Furthermore, positive regulation of vasculature development was completely absent in all cell states in the no suture group and was clearly enriched in many cell states after suturing. Although several cell states changed following duloxetine treatment, the enrichment for these biological processes remained in cell states in the treatment group, consistent with neovascularization not being prevented (Figure 1E).

## Discussion

In our study we leveraged single-cell RNA sequencing to describe transcriptomic signatures of cell populations involved in neovascularization of the cornea and performed a comprehensive molecular characterization of this pathological process – a hallmark of LSCD. We found a dramatic transformation of the original LSC population following suturing, resulting in the emergence of a large population of LSC-like cells (34%), compared to the no suture group (4% LSC). LSC-like cells in the injured corneas exhibited significantly lower expression of Pax6 and Trp63, consistent with previous reports identifying these as LSC markers in murine models and noting significant alterations under pathological conditions ^54–56^. Loss of *Trp63* (TP63 in humans) in particular has been significantly linked to pathological conditions such as corneal opacity ^57^, while patients transplanted with P63 protein-enriched LSCs demonstrate significantly higher success rates ^55^, underscoring the dual importance of P63 as a functional regulator and a prognostic marker in corneal regeneration. Some loss of ‘stemness’ relative to quiescent LSCs was therefore apparent in the present model, with inflammation likely causing these changes. For example, it was recently shown that inflammation reduces PAX6 expression in the murine cornea ^23^, corroborating its reduction in the present study after suturing. Additionally, reduced expression of *Krt5* and *Krt19* was detected in LSC-like cells in the suture group. In rodents, *Krt5* and *Krt19* have been reported as specific LSC markers ^54^. In mice, Krt5/Krt14 are co-expressed in LSCs as well as during differentiation to transient activating cells (TACs) together with Krt5/Krt12 ^54^ , whereas human corneas express K3/K12 ^58^. Krt19 has been reported to be normally expressed in murine limbal epithelium but not in the central corneal epithelium ^54^, emphasizing greater stem cell marker attenuation under suture conditions. Furthermore, the LSC-like cell state split into two states, which we annotated as ‘LSC-like 1’ and ‘LSC-like 2’, with distinct and opposing gene expression patterns of Sox9 and Cdk8, while retaining the limbal marker Gpha2. Sox9 has previously been shown to be localized to LSCs with a role in epithelial proliferation and differentiation in the context of wound healing ^59^. Its high and low expression in the LSC-like 1 and 2 populations, respectively, may indicate that the LSC-like 1 cells initiate symmetric cell divisions to maintain self-renewal of LSCs while LSC-like 2 cells give rise to asymmetric divisions leading to epithelial cell differentiation or transformation ^60^. Neither state, however, matches the original LSC state in terms of biological pathway/process enrichment. Tissue staining revealed an expanded GPHA2 expression pattern in both epithelial and stromal cells in sutured corneas, indicating that these cells have a altered cell state. This transformation could represent a trans-differentiation of these LSC-like cells or even cell de-differentiation, as was recently shown to occur in the murine cornea following limbal injury ^61^.

Duloxetine (Cymbalta) is known as a selective serotonin-norepinephrine reuptake inhibitor (SSNRI), used as an antidepressant and for treatment of other conditions such as major depressive disorder, general anxiety, and neuropathic pain. Its primary mechanism of action involves the modulation of neurotransmitters in the brain ^62^. Multiple lines of evidence support the anti-inflammatory effects of duloxetine with its treatment attenuating proinflammatory cytokines and various immune cells ^23,63–65^. Interestingly, duloxetine treatment in this study resulted in almost double the proportion of LSC-like 1 cells with high Sox9 expression among LSC-like cells (from 21% to 39%). Considering LSC-like 1 and 2 states, treatment enhanced their proportion from 34% to 53%. Expansion of the LSC-like 1 cell state corresponded to the potential rescue of PAX6 and GPHA2 expression at the protein level, highlighting the potential of duloxetine to restore LSC capacity following suturing. At the pathway level, TNF and MAPK signaling, axon guidance and negative regulation of epithelial cell migration were activated in LSCs of the normal non-sutured cornea. Following suturing, TNF, MAPK and axon guidance pathway enrichment were maintained in the LSC-like 1 cell state, while a switch was observed towards positive regulation of cell migration, cell adhesion, PI3K-Akt activation and positive regulation of inflammatory response. Interestingly, duloxetine treatment induced broad suppression of TNF, MAPK, PI3K-Akt, and suppressed the positive regulation of inflammatory response signaling, cellular adhesion and migration, while restoring the negative regulation of epithelial cell migration to a level observed in normal LSCs. This was corroborated at the protein level by strong basal and limbal epithelial suppression of TNFR2 encoded by *Tnfrsf1b* with duloxetine treatment. Supporting this finding, in a murine model of LPS-induced inflammation and depression-like conditions, it was previously demonstrated that duloxetine pre-treatment reduced serum TNFα levels ^64^. In summary, duloxetine induced a switch of pathway expression profile towards the uninjured corneal state in the LSC-like 1 population, a population that was simultaneously increased by duloxetine treatment, indicating a strong stem cell-mediated homeostatic effect of the drug.

Suturing also dramatically impacted basal epithelial cells, which declined by 75%, while basal cycling cells were fully suppressed. This indicates disruption of the normal basal renewal process, while the concomitant increase in LSC-like cells we hypothesize indicates a transformation of quiescent LSCs into an LSC-like state and/or a de-differentiation of basal cells into an LSC-like state. This was corroborated by broad expression of GPHA2 across all epithelial layers and even in stromal cells, indicating plasticity of a larger corneal cell population. These cell state changes may reflect a strong wound healing and repair response mediated by stem cell-like processes not limited to the corneal epithelium. Suturing also strikingly eliminated the differentiated non-basal epithelial cell state that originally comprised 36% of cells in the non-sutured cornea. A corresponding increase in conjunctival and conjunctival-like cells expressing keratins 13 and 15 suggests the transformation of differentiated superficial corneal epithelial layers into a non-transparent conjunctival phenotype, consistent with MUC5AC staining and the phenotypic opaque cornea observed clinically. Liang et al. previously established Krt13 as a reliable marker for LSCD, with its high expression indicating the presence of conjunctival epithelial cells on the corneal surface, a hallmark of LSCD ^66^. Note, however that this hypothesized transformation of non-basal epithelial cells to a conjunctival state does not result in a normal conjunctival state (cluster 8-9, 16-17) but instead modified conjunctival cells with low expression of *Muc4* and *Muc20* (cluster 18-20) and immune-like conjunctival cells expressing *S100a8/9*, *Ptprc* and *Cd52* (cluster 21).

In this context, duloxetine treatment completely suppressed these newly emergent conjunctival cell states after suturing (clusters 18-20 and 21), including *S100a8/9* suppression in conjunctival-like cells, mirroring previously reported *S100a8/9* suppression by duloxetine in an LPS-induced murine corneal inflammation model ^23^. Moreover, MAPK signaling, the positive regulation of vascular development and angiogenesis were suppressed by duloxetine. Previously, duloxetine was shown to suppress MAPK (MEK-ERK) signaling *in vitro* to activate PAX6 expression ^29^, which we now confirm *in vivo* at the transcriptomic level in this study. Additionally, MAPK downregulation was shown to not only maintain the self-renewal capacity of limbal stem cells but to prevent terminal differentiation of corneal epithelial cells ^67^, in agreement with our findings. Likewise, in a rat model of severe dry eye, MAPK inhibition preserved LSC self-renewal, underscoring the importance of this pathway in addressing epithelial dysfunction ^68^, while MAPK inhibition using MEK-inhibitor drugs restored corneal epithelial phenotype and marker expression in a mouse model of aniridia-associated keratopathy ^27^. Additionally, post-duloxetine treatment, CD86-expressing dendritic cells and IL-1β-expressing macrophages exhibited significantly diminished inflammation as evidenced by IL-1β staining. This reduction was evident in both limbal and central epithelial cells, suggesting partial suppression of inflammation. In a previous LPS-induced murine corneal inflammation model, duloxetine suppressed IL-1β as well as macrophage markers *Cd14* and *Cd52* ^23^.

Besides a reduction in cytokine-expressing dendritic cells and macrophages, duloxetine fully suppressed lymphocytes/NK cells while reducing the proportion of vascular endothelial cells and vessel-stabilizing mural cells and/or (myo)fibroblasts. The proportional reduction in these latter populations was modest, explaining the persistence of vessels and opacity in the cornea observed macroscopically. Given the multiple anti-inflammatory and antiangiogenic activities of duloxetine observed in the present suture model, more complete suppression of vascular and fibroblastic cell populations may be possible with a longer treatment period or alternatively, an optimized drug concentration. These findings substantiate previous reports suggesting that inflammatory and angiogenic processes in mouse corneal neovascularization models are inherently robust and resistant to short-term therapeutic interventions ^10^. Nevertheless, the impact of duloxetine on cell states and expression patterns in the present study was profound. Duloxetine treatment additionally uniquely triggered a stromal cell state with high Decorin expression throughout the stroma, revealed by gene expression and immunostaining. Decorin, a key extracellular proteoglycan predominantly expressed in the corneal stroma by stromal keratocytes and fibroblasts (Arts et al., 2024), plays a critical role in stromal fibrillogenesis by regulating collagen alignment, essential for maintaining corneal transparency and mechanical strength ^69^. This result suggests a duloxetine-enhanced stromal cell response to maintain transparency. Known for its anti-scarring properties, Decorin additionally acts as an antagonist of TGF-β and VEGFA, modulating ECM remodeling during wound healing ^69^. In alignment with this, angiogenic and vascular signaling reduction was noted in LSC-like cells following duloxetine treatment, although this did not rescue the neovascularization. Additionally, in the corneal stroma, Thy1, a mural cell and fibroblast marker, was absent in the normal corneal stroma but was highly expressed in neovascularized corneas. Known to be upregulated in vascular injury, Thy1 facilitates vascular remodeling ^46,70^. Duloxetine suppressed Thy1 gene and protein expression in the corneal stroma, along with a modest reduction in mural/fibroblast and vascular endothelial cell states, that was however insufficient to reduce neovascularization at the phenotypic level.

The enrichment of axon guidance pathways in non-epithelial cells and the emergence of Schwann cells following suturing is likely a response to the sub-basal and stromal nerve degeneration observed in the tissue by β III tubulin immunostaining and potentially explaining our observations on sub basal nerve degeneration and their possible activation following duloxetine treatment. Nerve regeneration involves a coordinated series of events that can become coupled with vascularization, involving endothelial cell migration, Schwann cell migration and axon guidance ^71,72^. The Schwann cell state was not fully diminished and axon guidance processes remained unaffected by duloxetine treatment. The nerve deficit after suturing may have partially accounted for the inability of the injured cornea to support LSCs, even with duloxetine treatment. The loss of corneal nerves following suture-induced neovascularization is also consistent with emergence of LSCD, as the limbal niche requires neurotrophic factors for its maintenance ^73,74^. Clinical findings of severely diminished corneal nerves in cases of congenital aniridia with LSCD and vascularized corneas illustrates this, with the degree of nerve loss correlating with severity of the LSCD and vascularization ^24,75^. Moreover, corneal nerve degeneration has been reported to be strongly associated with inflammation and an increase in dendritic cell density in various ocular ^76^ and systemic^77^ conditions.

Suturing also enriched PI3K-Akt signaling, primarily in LSC-like 1 cells. PI3K-Akt signaling regulates the proinflammatory response and vascular abnormalities ^78^ and plays a key role in the activation of vascular endothelial cells during angiogenesis ^79^. Duloxetine treatment suppressed this pathway in LSC-like 1 cells, highlighting a potential role in mitigating aberrant neovascularization.

In conclusion, we provide a comprehensive analysis of corneal neovascularization at the single-cell transcriptomic level, highlighting its overwhelming effects on epithelial cell states. With stimulation of inflammation and neovascularization, cells in the limbus and the basal corneal epithelium transform, resulting in the disappearance of homeostatic basal and differentiated epithelial cells and increase of limbal-like, conjunctival, inflammatory and vascular cell populations. This is likely the result of a wound healing and repair program whereby multiple cell types mount a coordinated response. Within this response, the potential exists for cell trans-differentiation, de-differentiation to more primitive states, as well as epithelial-to-mesenchymal transition; further investigation of temporal changes in cell states could yield important information in this regard. We found that Duloxetine initiates a program to restore corneal epithelial homeostasis while enhancing stem cell-like and stromal repair processes and simultaneously suppressing inflammation. The evidence indicates a promising therapeutic role for duloxetine in mitigating corneal inflammation and potentially neovascularization, although the optimal dose and duration of treatment require further investigation. Nevertheless, the detailed annotation of corneal cell states under normal and vascularized conditions in the murine cornea can provide insights into biomarkers and pathways of various cell types and serve as a basis for further study of the complex regulation of epithelial and LSCs and their broad roles in maintaining and repairing the cornea.

## Materials and Methods

### Animals and ethics

20 pathogen-free 6-8-week-old (postnatal day 42-56) male mice with C57BL/6 background were used, with animals bred and housed at the Linköping University animal facility, maintaining a standard dark-light cycle of 12:12 hours and providing access to food and water ad libitum. All animal experiments in this study were approved by the Linkoping Regional Animal Ethics Committee (Approval no. 10940-2021) prior to the commencement of the study and the procedures were performed in accordance with the Association for Research in Vision and Ophthalmology (ARVO) Guidelines for the Use of Animals in Ophthalmic and Vision Research.

### Preparation of duloxetine and vehicle Eye Drops

Duloxetine was obtained from European Pharmacopoeia (Duloxetine-hydrochloride, European Pharmacopoeia Reference Standard Y0001453). Duloxetine was prepared using duloxetine hydrochloride powder dissolved in PBS and the PH was adjusted to final neutral pH of 7.0-7.4. To investigate the toxicity profile of duloxetine, a range of duloxetine dosages (1, 10, 50, and 200 μM) were evaluated for the topical impact on the cornea, and 10 μM concentration (pH = 7.05, 21°C) was ultimately selected for further experimental validations ^23^. The solution was freshly prepared every three days and then sterilely distributed into eyedropper vials to be applied to mice corneas. Each time before using the eye drops, the solution was visually inspected to confirm the absence of particles or visual changes in colour during the study period.

### Mouse model of suture-induced inflammatory corneal neovascularization

Mice were first deeply anesthetized using intraperitoneal injection of ketamine (75 mg/kg) and xylazine (0.5 mg/kg). Additionally, topical anesthesia was applied using tetracaine hydrochloride eye drops. Three 10-0 nylon sutures (Serag Wiesner, Naila, Germany) were placed intrastromally in the cornea of mice in a figure-eight pattern ^6^, with each being separated by 120 degrees of the corneal circumference. Sutures were maintained in position for a period of 10 days to stimulate inflammation and corneal neovascularization., Mice were assigned into one of three experimental groups: Mice with corneas remaining un-sutured (hereafter called the ‘no suture’ group), mice with corneas sutured and receiving vehicle (PBS) eye drops (‘suture’ group) or 10 µM duloxetine (‘treatment’ group) administered topically twice daily for 10 consecutive days.

### Slip lamp imaging

Pupils were dilated using 0.5% (5 mg/ml) tropicamide and *in vivo* corneal examination was performed with the Micron III rodent slit lamp camera (Phoenix Research Laboratories, USA). Corneal transparency and blood vessels were visualized and photographed for later analysis.

### *In vivo* confocal microscopy imaging

Using laser-scanning *in vivo* confocal microscopy (HRT3-RCM; Heidelberg Engineering, Heidelberg, Germany) and while under general anesthesia, mouse corneas were examined at 10 days following suture placement and the images acquired were analyzed to assess inflammation status. The microscope provided an en-face view of a 400 × 400 μm central corneal area at a selectable depth. for statistical analysis between groups, three stromal depths were chosen with 6 images captured at each depth.

### Whole-mount corneal immunostaining and fluorescent imaging

Mice were euthanized on day 10 under deep general anesthesia and the entire corneas with scleral rims were isolated. The full thickness cornea was prepared and dedicated for flat mounting and processed for wholemount staining. Corneas were fixed in 1.3% PFA for one hour, then rinsed in PBS three times for 5 minutes. Next, corneal tissues were blocked with 1% normal goat serum in 0.1% Triton-X in PBS for 1 hour at room temperature and double-stained with antibodies against the neuronal-specific marker beta-III-tubulin (1:200 abcam 78078), and vascular endothelial marker CD31 (PECAM-1) (1:50, Abcam 28364). For this purpose, corneas were incubated overnight in primary antibodies and after a subsequent wash in washing buffer (0.3% Tween in PBS), incubated for 48 hours with secondary anti-mouse (Alexa-Fluor 488) antibodies and thereafter overnight with anti-rabbit (Alexa-Fluor 647) antibody at 4°C. Finally, corneas were washed and mounted in slowfade Diamond Antifade Mountant (Invitrogen, S36972). Blood vessel sprouts and total cornea neurons were imaged using a laser-scanning confocal fluorescence microscope (Zeiss LSM 800) with a 10xobjective lens. Using Image J software (Image J, NIH, USA) ^80^, the area positive for CD31, indicating blood vessels, was quantified and normalized to the total corneal area, yielding the percentage coverage. Furthermore, Image J was employed to quantify corneal nerves in the vehicle and duloxetine groups. Changes in beta III tubulin-positive subbasal and stromal nerves were evaluated in terms of their proportional coverage of the total corneal wholemount area expressed as percentages. For both analyses, the circumferential limbal vessel arcade and nerves were used as the outer boundary to define the total corneal area.

For immunofluorescence staining, PFA-fixed-paraffin-embedded tissues were processed into 5μm thick sections. Sections were subsequently deparaffinized and rehydrated following a standard protocol of consecutive 5 min immersions in containers with xylene (twice) and descending ethanol concentrations (2x 100%, 95%, 80%, 70%) and finally immersed in tap water. Antigen retrieval was performed using the DAKO PT LINK 200 antigen unmasking system with acidic buffer and blocked with 1% BSA, 5% normal goat serum in 0.3% Triton-X in PBS for 1 hour. Primary and secondary antibodies were diluted in 1% BSA 5% goat serum 0.1% Triton-X in PBS. Overnight incubation at 4°C with primary antibodies was performed in a hydrated chamber. The antibodies used were against MUC5AC (1:50, clone 45M1, Invitrogen MA512178 by Fiscer-sci, product code 11394583), GPHA2 (1:50, clone G-3, Santa Cruz sc-390194), Thy1 (1:100, Invitrogen by Thermo-Fisher HL1766, catalog # MA5-47174), CD86 - B7-2 (1:100, Invitrogen by Thermo-Fisher, catalog # 14-0862-82), Decorin (1:100, Proteintech by Thermo-Fisher, catalog # 14667-1-AP), TNFR2 (1:100, Invitrogen JM113-01 by Thermo-Fisher, catalog # MA5-32618), IL-1 beta (1:200, Invitrogen by Thermo-Fisher, catalog # P420B), and PAX6 (1:100, Abcam EPR15858). After the overnight incubation, section slides were washed with PBS 1% Tween-20 and then the secondary antibody was applied for 2 hours at ambient temperature. Secondary antibodies were goat anti-Rabbit IgG-Alexa Fluor™ Plus 647 (1:400, Invitrogen by Thermo-Fisher, catalog # A32733), Goat anti-Mouse IgG-Alexa Fluor™ Plus 488 (1:400, Invitrogen by Thermo-Fisher, catalog # A32723), and Goat anti-Rat IgG-Alexa Fluor™ 546 (1:400, Invitrogen by Thermo-Fisher, catalog # A-11081). Sections were washed with PBS 0,1% Tween-20 and a mounting medium with DAPI (SlowFade™ Diamond Antifade Mountant with DAPI, Molecular Probes™ S36964 by Fisher-sci, product code 15451244) was applied before coversliping. Images were taken under an upright Zeiss LSM800 confocal microscope.

### Statistical analysis

The vessel coverage and loss of subbasal and stromal nerves were compared across no suture, suture and treatment groups. All data were statistically analyzed and graphed using GraphPad Prism software (Version 9.2.0, GraphPad Software, San Diego, CA, USA). The results in column plots are presented as mean ± SD from given sample sizes (n). Comparisons were made using Student’s t-test for unpaired data or one-way ANOVA followed by Bonferroni ‘s post-hoc test for multiple comparisons. A significance level of P < 0.05 was considered as statistically significant.

### Single-cell preparation from mouse cornea and sequencing

Mouse corneas were dissected from harvested eyes and digested in Dispase II (2.4 IU/ml) at 4°C overnight with an additional incubation for about 2 hours at 37 °C with collagenase type I (Gibco, 1 mg/ml) in Eagle’s Minimum Essential Medium (EMEM) containing 10% FBS. Isolated cells were pelleted by centrifugation and further digested into single cells with 0.25% trypsin in EDTA (1mM) for 20 minutes at 37°C. During this time, cell suspension was gently pipetted intermittently to produce single cells. Then the cells were washed (centrifuged-resuspended after discarding supernatant) once with EMEM with 10% FBS to deactivate and remove trypsin and two times with sterile PBS containing 0.04% BSA (Ca2+/Mg2+ free). Finally, cells were resuspended in 30-40 μl PBS containing 0.04% BSA and counted to achieve a number within the range of 700 to 1200 cells/μl.

For scRNA-Seq, cells were captured and libraries generated using the Chromium Single Cell 3′ Library & Gel Bead Kit, version 3 (10x Genomics). scRNA-Seq libraries were sequenced to 50,000 reads per cell on an Illumina NovaSeq 6000.

### Pre-processing and quality control of scRNA-seq

Cellranger count was run with Cellranger 7.0.1 ^81^ with default parameters and with mm10 to retrieve the matrix, barcodes and features files necessary for downstream analysis. scRNA-seq datasets were analyzed in Python with Scanpy version 1.9.6 ^82^, as described on GitHub: https://github.com/Arts-of-coding/Perturbation-of-epithelial-and-limbal-stem-cell-identity-in-pathologic-corneal-neovascularization. scRNA-seq cells were selected with a minimum count of 2000, a maximum count of 100.000, a feature number higher than 1000 and lower than 8000 as well as a mitochondrial percentage lower than 30%. Estimated doublets were removed with Scrublet 0.2.3 ^83^.

### Clustering and integration of scRNA-seq datasets

Clustering on scRNA-seq data on the three conditions (no suture, suture and treatment) was performed with Leiden clustering ^31^ using 30 dimensions and a neighbors parameter 10 for to identify cell clusters. Cell states were defined by combining clusters sharing expression of a multitude of marker genes. The most optimal clustering resolutions were determined with Clustree version 0.5.0 ^84^ and by analyzing the clustering scores for the silhouette index ^85^, Davies-Bouldin index ^86^ and Calinski-Harabasz index ^85^, resulting in final clustering resolutions of 0.5, 0.65, and 0.5 for the three conditions, respectively. The three datasets were first annotated based on marker genes individually and were then integrated with scVI ^32^ (scvi-tools version 1.0.0). The parameters for the Variational Autoencoder were selected as 2 for “n_layers”, 30 for “n_latent” and “nb” for “gene_likelihood”.

### GO-term enrichment and KEGG pathway analysis

Gene ontology enrichment ^87^ and KEGG pathway enrichment ^52^ were performed on cell states in the combined data from all three groups (no suture, suture, treatment). Gene set enrichment was achieved using the databases: GO Biological Process 2021 from Mouse and KEGG 2019 Mouse in decoupler-py 1.1.1 ^88^ by using the highly variable genes within the combined dataset (Table S2). From the enriched pathways, the following KEGG terms previously suggested to be involved in neovascularization, inflammation or duloxetine mode of action: were further investigated: Chemokine signaling pathway, TNF signaling pathway, MAPK signaling pathway, PI3K-Akt signaling pathway, Leukocyte transendothelial migration, ECM-receptor interaction and Axon Guidance ^10,50,89^. Similarly, the GO identifiers: GO:0045596, GO:0045785, GO:0050729, GO:0010633, GO:1900117, GO:0010811, GO:0030335, GO:1904018, GO:0045766, GO:0071354, GO:0070102 were pre-selected. These terms were selected with an adjusted (Benjamini-Yuketeili) p-value of <0.05 ^90^.

### Data and code availability

Datasets containing raw scRNA-seq data used in this study are available under the Gene Expression Omnibus (GEO): <u>GSE294213</u>. Generated single cell objects are available on Zenodo at https://doi.org/10.5281/zenodo.15211589. The processing workflow and documentation of code used is available on GitHub: https://github.com/Arts-of-coding/Perturbation-of-epithelial-and-limbal-stem-cell-identity-in-pathologic-corneal-neovascularization. GRCm38 (mm10) was downloaded from 10X Genomics and was used for mapping reads to the genome. A web-based application was developed to visualize scRNA-seq datasets interactively. The web application is available at: https://huggingface.co/spaces/Zhou-group/mousesuture.

## Data availability statement

The data supporting the findings of this study are available from the corresponding author upon request.

## Acknowledgments

Funding for this study was obtained (to N.L. and H.Z.) from the European Joint Programme on Rare Diseases (EJP-RD) under the project AAK-INSIGHT, Grant. No. EJPRD20-135.

## Author Contributions

M.A: Conceptualization, performed experiments and data collection, data analysis and interpretation, manuscript writing and review. J.A.A: data analysis and interpretation, manuscript writing and review. D.J: performed experiments and data collection, manuscript writing and review PM: performed experiments and data collection, manuscript writing and review. A.D: performed experiments and data collection, manuscript writing and review D.A: Conceptualization, manuscript writing and review H.Z: coordination of data analysis, funding, data interpretation, manuscript writing and critical review. NL: Conceptualization, study design, coordination of experiments, funding, data interpretation, critical manuscript review and revisions.

## Declaration of interest

The authors have no conflicts of interest to declare.

## Declaration of Generative AI and AI-assisted technologies in the writing process

The authors confirm that no generative AI or AI-assisted technologies were used in the writing or editing of this manuscript.

## Notes

### Competing Interest Statement

The authors have declared no competing interest.

