## Supplemental material for "Perturbation of epithelial and limbal stem cell identity in a mouse model of pathologic corneal neovascularization"

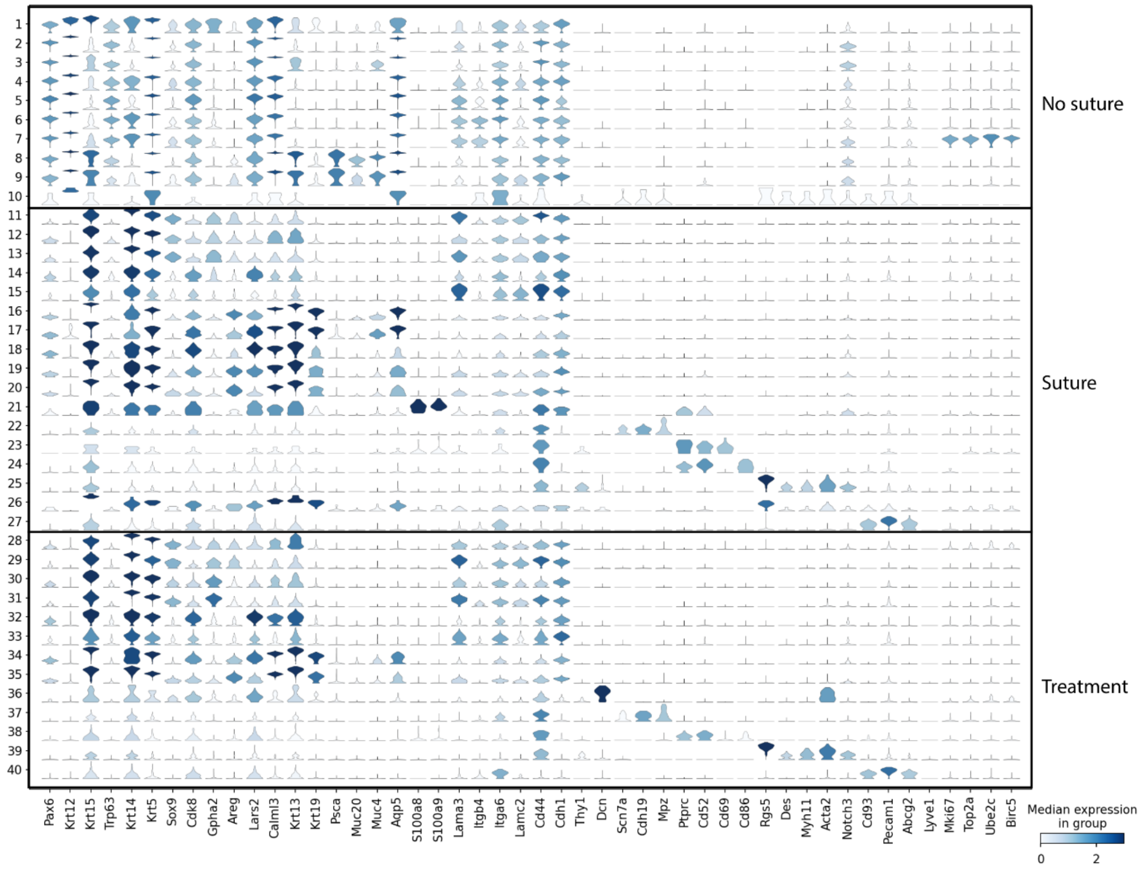


**Supplementary Figure 1:** Violin plots of marker gene expression of unbiased clusters across no suture, suture and treatment groups.

**Supplementary Table 1:** Proportional distribution of various cell states based on experimental condition, as shown graphically and by color in Figure 2 (A-D). Within each condition, 100% of cells in the cornea were distributed across the named cell states.

| Condition | Cell state | Proportion of cells/condition (%) |
|---|---|---|
| No suture | 1: LSC | 4.30 |
| | 2-3: Non-basal epithelial | 36.17 |
| | 4-6: Basal epithelial | 35.34 |

| | | |
|---|---|---|
| | 7: Basal cycling | 5.96 |
| | 8-9: Conjunctiva | 17.96 |
| | 10: Non-epithelial | 0.27 |
| Suture | 11-13: LSC-like 1 | 20.54 |
| | 14: LSC-like 2 | 13.64 |
| | 15: Basal epithelial | 7.61 |
| | 16-17: Conjunctiva | 15.58 |
| | 18-20: Conjunctiva_Muc4/Muc20-low | 25.18 |
| | 21: Conjunctiva/immune-like | 0.81 |
| | 22: Schwann cells | 4.51 |
| | 23: Lymphocytes/natural killer cells | 0.37 |
| | 24: Dendritic cells/macrophages | 1.65 |
| | 25: Mural cells | 6.69 |
| | 26: Vascular/conjunctiva-like | 0.47 |
| | 27: Vascular endothelial cells | 2.96 |
| Treatment | 28-31: LSC-like 1 | 38.64 |
| | 32: LSC-like 2 | 14.89 |
| | 33: Basal epithelial | 8.15 |
| | 34-35: Conjunctiva | 26.04 |
| | 36: Stromal cells | 2.27 |
| | 37: Schwann cells | 2.65 |
| | 38: Dendritic cells/macrophages | 2.01 |
| | 39: Mural cells | 3.34 |
| | 40: Vascular endothelial cells | 2.01 |

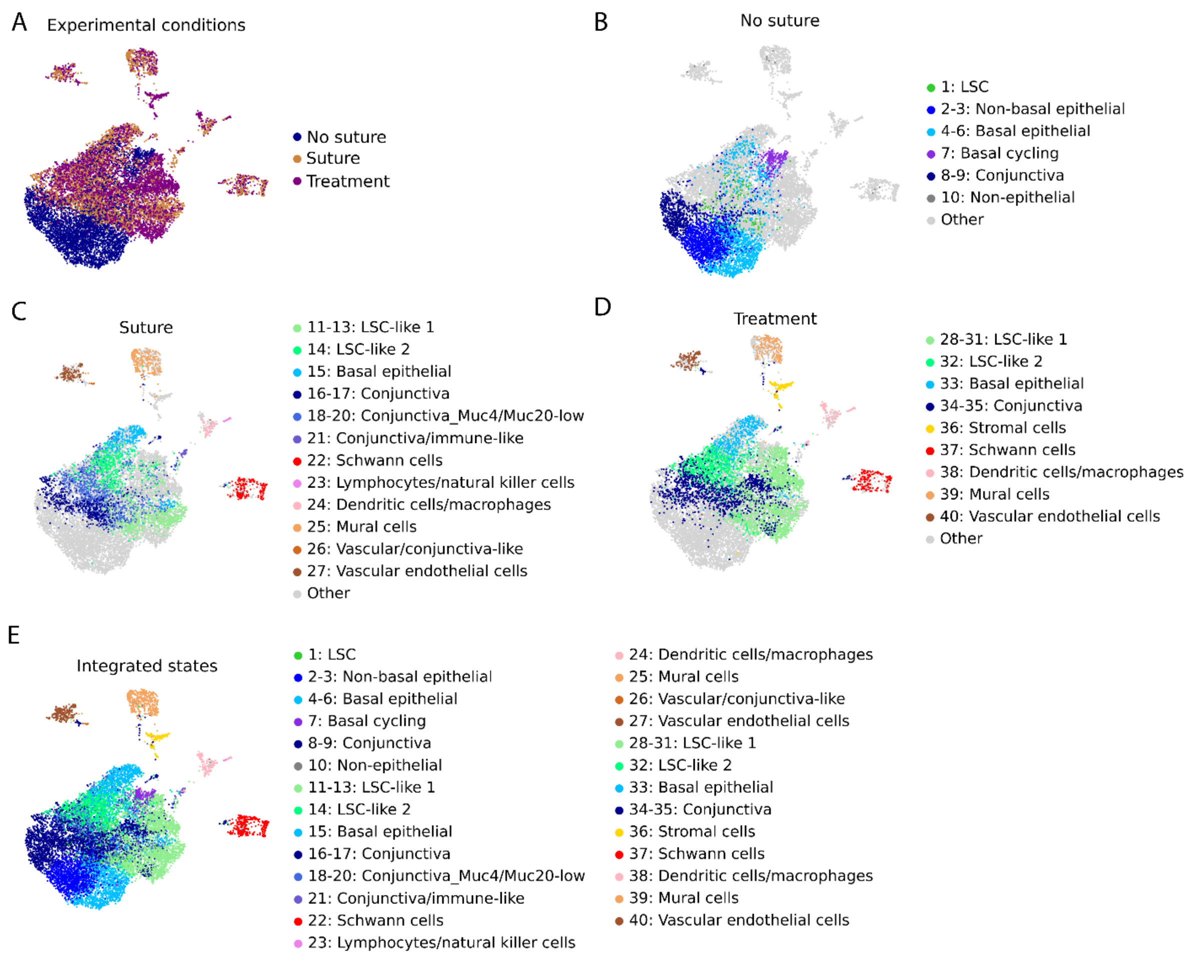


**Supplementary Figure 2: Dataset integration of cell states in no suture, suture and duloxetine treated corneas.** (A-E) Integrated UMAP of experimental groups (A), cell states identified in no suture (B), cell states identified in suture (C), cell states identified in treatment (D) and all cell states integrated across the three experimental groups (E).

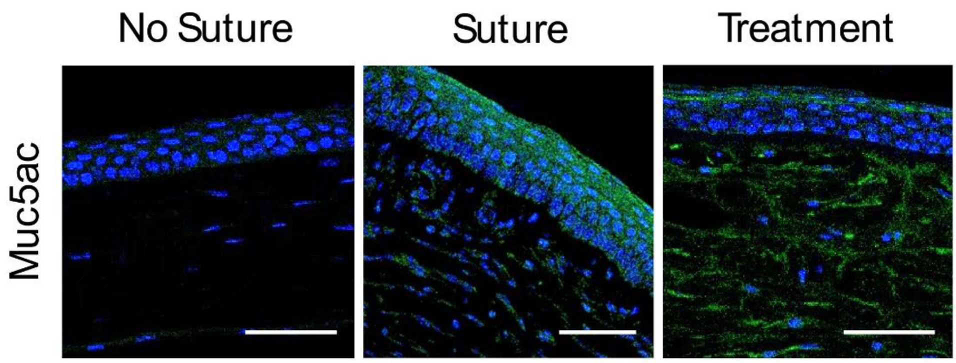

**Supplementary Figure 3: Staining of the conjunctiva marker Muc5ac.** While corneas in the no suture group did not express Muc5ac, its expression was detected in superficial layers of the corneal epithelium with suturing. Duloxetine treatment ameliorated the epithelial expression of this conjunctival marker, where it was apparent primarily in the most superficial epithelial layer.

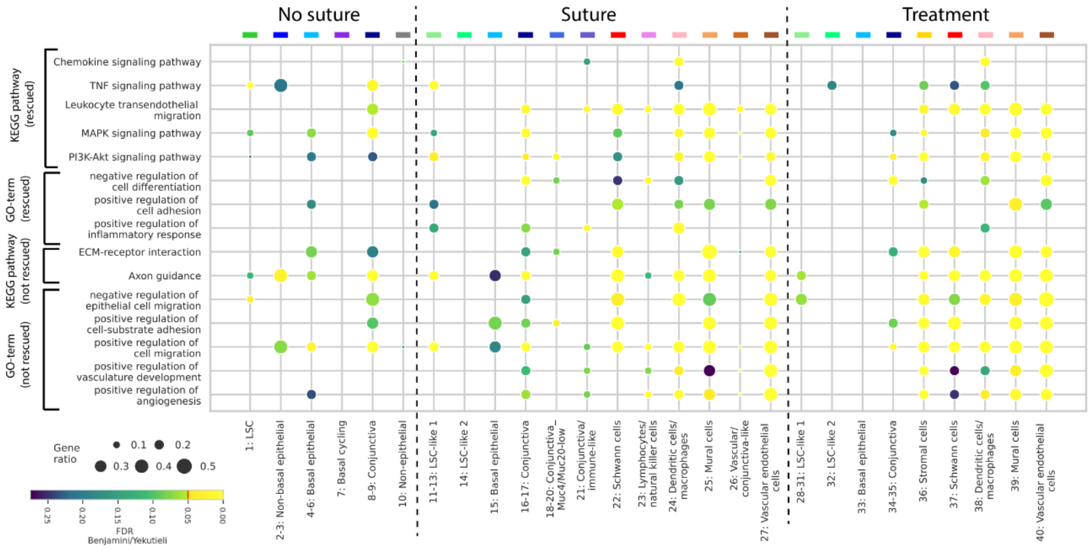


**Supplementary Figure 4**: **Gene set enrichment characterization of disease processes on the single-cell landscape of sutured corneas and effects by Duloxetine.** Preselected gene sets associated with suture-induced disease processes underwent GO-Term and KEGG pathway enrichment analysis across all cell states within the three experimental groups (no suture, suture and treatment).

The data are presented as the p-value adjusted using the Benjamini–Yekutieli method.
